# Cerebellar re-encoding of self-generated head movements

**DOI:** 10.1101/126086

**Authors:** Guillaume P. Dugué, Matthieu Tihy, Boris Gourévitch, Clément Léna

**Author notes:** Equal contributions (Guillaume P. Dugué, Matthieu Tihy).

## Abstract

Head movements are primarily sensed in a reference frame tied to the head, yet they are used to calculate self-orientation relative to the world. This requires to re-encode head kinematic signals into a reference frame anchored to earth-centered landmarks such as gravity, through computations whose neuronal substrate remains to be determined. Here, we studied the encoding of self-generated head movements in the caudal cerebellar vermis, an area essential for graviceptive functions. We found that, contrarily to peripheral vestibular inputs, most Purkinje cells exhibited a mixed sensitivity to head rotational and gravitational information and were differentially modulated by active and passive movements. In a subpopulation of cells, this mixed sensitivity underlay a tuning to rotations about an axis defined relative to gravity. Therefore, we show that the caudal vermis hosts a re-encoded, gravitationally-polarized representation of self-generated head kinematics.

## Introduction

Self-orientation is largely dependent on modalities that document head movements, including neck proprioception, optic flow and, most notably, vestibular inputs. Vestibular signals are essential for stabilizing gaze (du Lac et al., 1995) and for computing head direction, spatial maps and navigation trajectories (Stackman et al., 2002; Yoder & Taube, 2014; Wallace et al., 2002). These signals originate from two categories of skull-anchored inertial sensors: gyroscope-like structures (semi-circular canals), which transduce head angular velocity, and accelerometer-like structures (otolith organs), which are activated indifferently by accelerated linear motion and by gravity. Gravity provides an absolute directional cue on the external world and is effectively derived from vestibular inputs in the vestibular system (Merfeld et al., 1999; Angelaki et al., 1999), allowing the brain to align the axes of eye rotations with the direction of gravity (Hess, 2003). Head direction cells are also anchored to a reference frame aligned with gravity rather than to the animal’s locomotor plane (Taube et al., 2013; Finkelstein et al., 2016; Wilson et al., 2016; Olson et al., 2016); their activity, which relies on the temporal integration of head angular velocity signals (Song & Wang, 2005), thus also requires information on head orientation relative to gravity (head tilt) (Yoder & Taube, 2009). Indeed, identical activations of semi-circular canals may affect the azimuth and elevation of the head in very different ways depending on head tilt: for example a rotation about the interaural axis will lead opposite changes of azimuth if the head is tilted with the left or right ear down. Therefore, understanding how the brain computes head direction requires to identify the neuronal substrate of the operations transforming skull-bound angular velocity into changes of azimuth and elevation.

Lesion data suggest that the caudal cerebellar vermis, a brain region receiving multimodal sensory cues related to head kinematics and orientation (Quy et al., 2011; Yakusheva et al., 2013), plays a pivotal role in the discrimination of gravity (e.g. Tarnutzer et al., 2015). Moreover, a distinct population of caudal cerebellar Purkinje cells in monkeys dynamically reports head tilt during passive whole-body movements (Yakusheva et al., 2007; Laurens et al., 2013). We therefore hypothesized that this structure might also host also a representation of head rotations anchored to the direction of gravity.

A considerable literature has described the responses of caudal vermis Purkinje cells to passively experienced head movements (Barmack & Yakhnitsa, 2011). However, these movements only covered the lower range of frequencies and amplitudes observed during active self-motion (Carriot et al., 2015). Moreover, despite the remarkable linearity of early vestibular information processing (Bagnall et al., 2008), the high amplitude of active movements might recruit vestibular afferents in a non-linear way (Schneider et al., 2015). In addition, studies in mice and monkeys have revealed that active and passive head movements are processed in fundamentally different ways within the vestibular nuclei (Cullen & Roy, 2004), which are highly interconnected with the caudal cerebellar vermis. Thus, the principles of vestibular coding in passive conditions might not apply to the active condition. We therefore decided to study the encoding of head movements in the caudal cerebellar vermis in freely moving rats, while monitoring the movements of their head using a miniature inertial sensor.

## Results

### Kinematics of self-generated head movements

Combined recordings of cerebellar activity and head movements were obtained in 16 freely-moving rats (Fig. 1A). In average, head rotations occurred more frequently and swiftly along the pitch and yaw axes than along the roll axis (Figure 1–figure supplement 1A,B), with typical angular speed in the 18−287 °/s range (average 2.5−97.5% percentiles calculated for velocities > 15°/s, *n* = 16 rats). Angular velocity (**Ω**) signals displayed multi-peaked power spectra spanning frequencies up to 20 Hz (Figure 1-figure supplement 1C) and showed strong temporal autocorrelation over a short timescale (< 0.2 s, Figure 1–figure supplement 2A). The gravitational (A^G^) and non-gravitational (A^nG^) components of acceleration (A) were calculated using an orientation filter algorithm (Madgwick et al., 2011, Figure 1–figure supplement 1D). A^G^ accounted for almost all (99%) of the power of A below 2 Hz, and for only 9%of it in the 2–20 Hz range (*n* = 16 rats, Figure 1–figure supplement 1E), indicating that the low-frequency component of acceleration (< 2Hz) mostly contained head tilt information. Consistent with this, A^G^ displayed temporal autocorrelation over long timescales (> 5s, Figure 1–figure supplement 2B). A^nG^ varied at the same timescale as **Ω** and exhibited the same amplitude and temporal correlation pattern with **Ω** as linear tangential acceleration predicted from head rotations (Figure 1–figure supplement 2C–E). This suggests that, in our conditions, A^nG^ signals arose primarily from head rotations.

**Figure 1.**
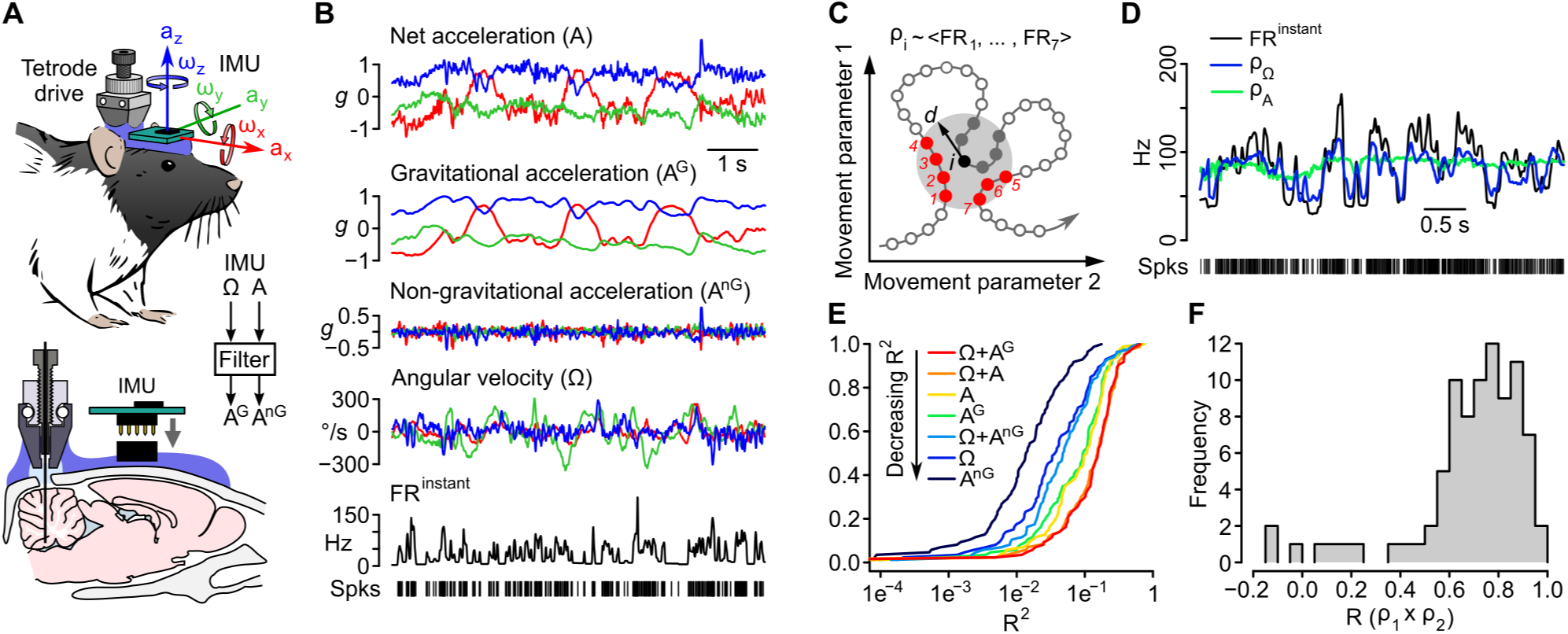
Caudal cerebellar units are sensitive to different combinations of rotational and gravitational information. (A) Orientation of the inertial measurement unit (IMU) on the animal’s head and tetrode placement. An algorithm (filter) was used to calculate the gravitational (A^G^) and non-gravitational (A^nG^) components of acceleration (A) using angular velocity information (**Ω**). (B) Traces showing the instantaneous firing rate (FR^instant^) of an example unit and inertial signals recorded simultaneously (A and **Ω**) or calculated offline (A^G^ and A^nG^). (C) Principle of the resampling method. Recordings of head movements can be described as sequences of points in a multidimensional parameter space (circles and line, here represented in a 2D space). At a given time point *i* (black circle) the estimated firing rate *ρ_i_* is the mean of FR^instant^ values observed for neighboring points in the parameter space within a distance *d* (red circles) that did not occur immediately before or after *i* (filled gray circles). (D) Firing rate estimates calculated using **Ω** (*ρ*_**Ω**_) or A (*ρ*_*A*_) and FR^instant^ of an example unit. (E) Cumulative distribution of *R*^2^ for firing rate estimates calculated based on different combinations of inertial parameters (*n* = 86 units). (F) Distribution of Pearson correlation coefficients between independent firing rate estimates (*n* = 86 units).

### Cerebellar units exhibit a mixed sensitivity to head angular velocity and gravitational acceleration

A total of 86 units were recorded (Fig. 1A and Figure 1–figure supplement 3A–C) and classified into putative Purkinje cells (90%), Golgi cells (5%) and mossy fibers (5%) according to established criteria (Figure 1–figure supplement 3E; van Dijck et al., 2013). Purkinje cells exhibited irregular inter-spike intervals (ISI) at rest (average CV: 0.95 ± 0.58, calculated for periods of immobility isolated from 76 cells), resulting in sharp fluctuations of the instantaneous firing rate even during immobility (Figure 1–figure supplement 3D). To infer the influence of head movements on recorded units, we designed a model-free resampling technique that calculates firing rate estimates based on the repeated occurrence of similar combinations of inertial parameters (Fig. 1C,D). The square of the Pearson correlation coefficient (*R*^2^) between estimated and observed firing rates was taken as a measure of the fraction of instantaneous firing rate fluctuations that could be explained by a given combination of inertial parameters. Overall, firing rate fluctuations were better explained when considering **Ω** combined with either A or A^G^ (mean *R^2^* = 0.17 ± 0.13 in both cases, *n* = 86 units;Fig. 1E). Considered alone, all inertial parameters (**Ω**, A, A^G^ or A^nG^) explained significantly smaller fractions of firing rate fluctuations than combinations of **Ω** and A or A^G^ (*p*< 5e−3, *n* = 86 units). *R*^2^ values were always greater for A^G^-based than for A^nG^-based estimates (*p* < 8e−9 for both A^G^ vs. A^nG^ and **Ω** + A^G^ vs. **Ω** + A^nG^, *n* = 86 units), showing that gravitational information dominated the effect of acceleration on firing rate. *R*^2^ values for estimates obtained with **Ω** or **Ω** + A^nG^ were similar (*p* = 0.20, *n* = 86 units), consistent with a redundancy of these parameters due to their coupling (Figure 1–figure supplement 2E). This analysis suggests that cerebellar units preferentially exhibited a mixed sensitivity to head rotations and head tilt.

The correlation between estimated and observed firing rates is intrinsically limited by the strong fluctuations of the latter. To assess the robustness of our method, we examined whether independent portions of the same recordings yielded consistent estimates (see Methods). We found indeed that in 80% of the units, the Pearson correlation coefficient (*R*) between independent estimates was greater than 0.6 (Fig. 1F), showing the ability of our method to consistently capture the link between head movements and firing rate modulations.

### The influence of head movements on firing rate is independent of visual cues but varies when movements are self-generated or passively experienced

The instantaneous firing rate of neighboring cells (recorded from the same site, Fig. 2A,B) displayed positive correlations (*R* = 0.14 ± 0.26, *n* = 35 pairs; *p* = 1.4e−3) that were lost if the spike train of one cell was time-reversed (*R* = 0.00 ± 0.03, *p* = 0.49; Fig. 2C). Correlations were higher (*p* = 6.9e−3) when comparing firing rate estimates (*R* = 0.31 ± 0.47, *p* = 1.3e−3; Fig. 2C,D), showing that neighboring cells tended to share similar sensitivities to head movements.

**Figure 2.**
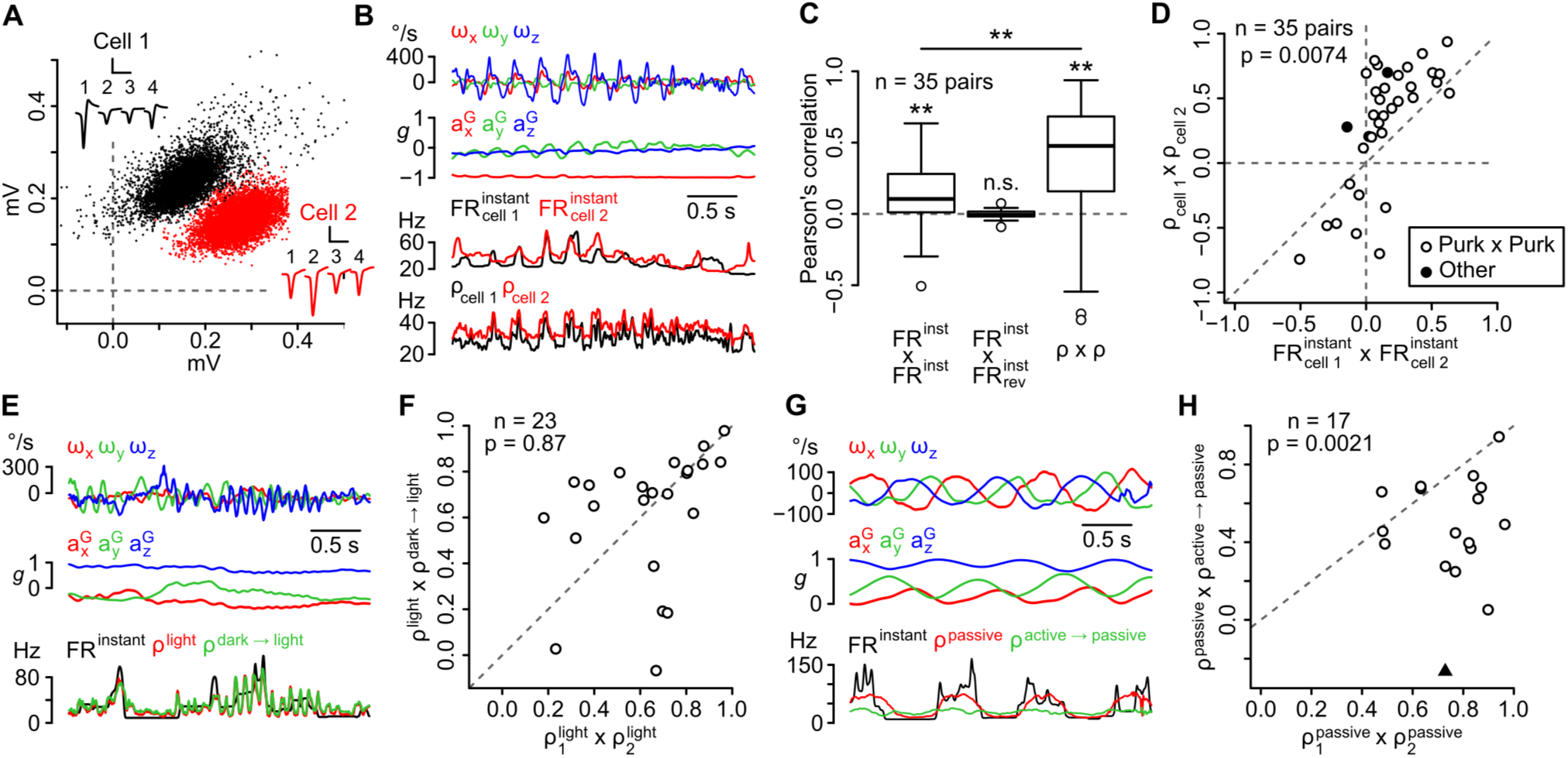
The sensitivity of recorded units is similar in the same recording site and does not depend on visual cues, but differs between active and passive movements. (A) Amplitude of sorted spikes on a pair of channels and average spike waveforms of two neighboring units (scale bars: 0.3 mV and 1 ms for cell 1,0.15 mV and 1 ms for cell 2). (B) Example traces showing inertial parameters and the instantaneous and estimated firing rates of the two cells shown in *A*. (C) Boxplots of Pearson correlation coefficients between instantaneous (FR^instant^ × FR^instant^) or estimated (*ρ* × *ρ*) firing rates of neighboring cells (*n* = 35 pairs, *\*\*p*< 0.01). The correlation between FR^instant^ was lost if the firing rate of one of cell was time-reversed (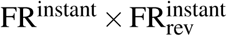, *p* = 0.49). (D) Graph comparing Pearson correlation coefficients between firing rate estimates (*ρ_cell1_* × *ρ_cell2_*) and instantaneous firing rates 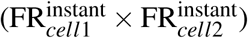 of neighboring cells (*n* = 35 pairs). Pairs of putative Purkinje cells (*n* = 32) are shown in white. (E) Example traces showing inertial parameters and FR^instant^ for one example unit recorded in the light block. Color traces are firing rate estimates for the same recording calculated using data from the same block (*ρ_light_*) or from the dark block (*ρ*^dark→light^). (F) Graph comparing Pearson correlation coefficients between independent firing rate estimates in the light block 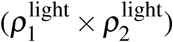 and between estimates of the firing rate in the light block calculated using data from either the light or dark block (*ρ*^light^ × *ρ*^dark→light^). All units corresponded to putative Purkinje cells (*n* = 25). (G) Example traces showing inertial parameters and FR^instant^ for one example unit recorded in the passive block. Color traces are firing rate estimates for the same recording calculated using data from the same block (*ρ_passive_*) or from the passive block (*ρ*^active→passive^). (H) Graph comparing Pearson correlation coefficients between independent firing rate estimates in the passive block 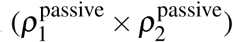 and between estimates of the firing rate in the passive block calculated using data from either the passive or active block (*ρ*^light^ × *ρ*^dark→light^). All units corresponded to putative Purkinje cells, except one classified as a Golgi cell (black triangle; *n* = 25 units in total).

We then examined whether changing experimental conditions affected the units’ sensitivity by comparing independent firing rate estimates obtained by resampling from the same or from different conditions (see Methods). Overall, the presence or absence of light did not change the correlation between independent estimates (*p* = 0.87, *n* = 23; Fig. 2E,F), showing a limited influence of visual cues on the units’ sensitivity to head movements. However, independent estimates differed significantly for active vs. passive movements (*p* = 2.1e−3, *n* = 17; Fig. 2G,H), indicating that in many cases the units’ coding schemes differed when movements were self-generated or passively experienced.

### Subsets of cerebellar units are specifically tuned to either rotational or gravitational information

Although most units exhibited a mixed gravitational and rotational sensitivity ( Fig. 1E), a fraction of them appeared to be mostly tuned to **Ω** or A^G^. We isolated 6 **Ω**-selective and 12 A^G^-selective units by picking units for which *R*^2^ values were at least 8 fold greater for one parameter than the other (Fig. 3A). The sensitivity of **Ω**-units was examined by computing inertio-temporal receptive fields (average instantaneous firing rate timecourse around specific angular velocity values). These plots showed sharp and bidirectional tuning to specific combinations of rotation axes (Fig. 3B). 3D plots representing the firing rate as a function of the three components of **Ω** displayed clear firing rate gradients along specific directions, showing that these units were tuned to specific 3D rotations (Fig. 3C) and Movie 1). Rotational sensitivity was described by a vector defined by the coefficients of the firing rate vs. **Ω** linear regression (Fig. 3D). The norm of this vector, in Hz/(°/s) or /° (reflecting the gain of the unit’s response), was maximal (0.28 ± 0.15 °^−1^, *n* = 6) at an optimal lag (0.5 ± 21.8 ms, *n* = 6), defining an optimal sensitivity vector **ω*_opt_*** whose direction indicated to the unit’s preferred rotation axis (Fig. 3C). **Ω**-units corresponded to putative mossy fibers (*n* = 2) and Purkinje cells (*n* = 4), and exhibited preferred rotation axes clustered around the excitatory direction of semi-circular canals (Fig. 3F). Units with significant **ω*_opt_*** vectors that were neither classified as **Ω**- nor as A^G^-units (*n* = 53) displayed smaller sensitivities (0.07 ± 0.05 °^−1^) and longer optimal lags (30 ± 142 ms), and corresponded in majority (96%) to Purkinje cells.

**Figure 3.**
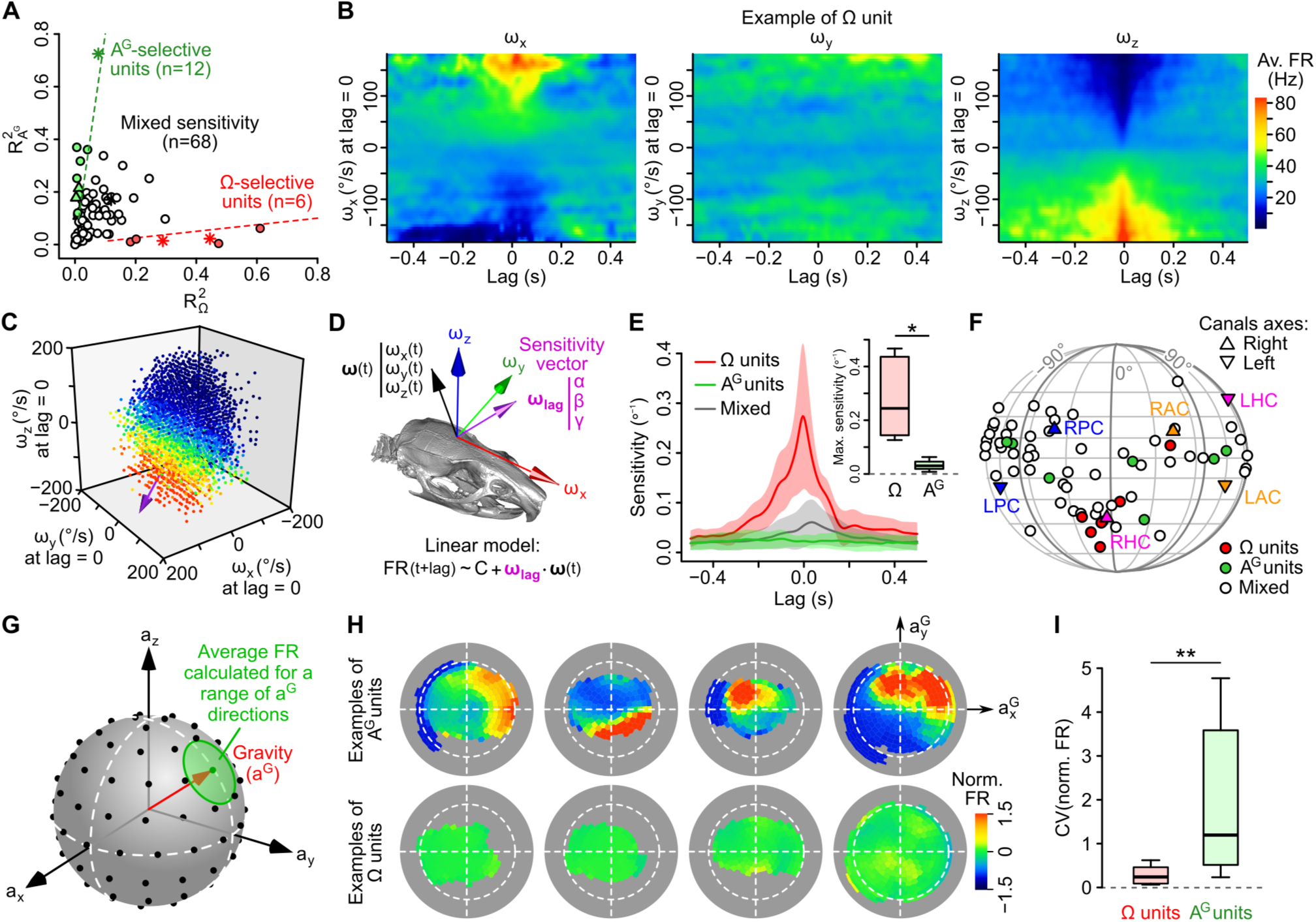
Subsets of caudal cerebellar units display preferential sensitivity to either head angular velocity or head tilt. (A) Comparison of *R*^2^ values calculated based on gravitational acceleration 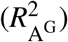 or angular velocity 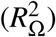 for all 86 units. Dashed lines delineate units with an *R*^2^ at least 8 times greater for one parameter than for the other and greater than 0.1 for the preferred parameter. Putative Purkinje cells, Golgi cells and mossy fibers are represented by empty circles, filled triangles and stars, respectively. (B) Inertio-temporal receptive fields of one example **Ω**-unit. (C) Firing rate (color-coded) of one **Ω**-unit (same as *B*) plotted as a function of the three components of angular velocity (see also Movie 1). The unit’s optimal sensitivity vector (calculated using linear regression) is represented in purple (arbitrary scale). (D) Linear model used to characterize the units’ rotational tuning. For a given lag, the model assumes a linear tuning of firing rate to a preferred sensitivity vector ***ω_lag_***. (E) Average (±*SD*) rotational sensitivity (norm of ***ω_lag_***.) plotted vs. lag for **Ω**-units (*n* = 6), A^G^-units (*n* = 7) and other units (*n* = 53). Only units with significant sensitivity were included. Inset:boxplot of the sensitivity of **Ω**- and A^G^-units at their optimal lag. **p = 1.2e−3. (F) Direction of optimal sensitivity vectors of **Ω**-units, A^G^-units and other units plotted on a pseudocylindrical projection. Triangles point up (resp. down) represent the excitatory direction of rotation of right (resp. left) semi-circular canals. LPC/RPC: left/right posterior canals; LHC/RHC: left/right horizontal canals; LAC/RAC: left/right anterior canals. (G) Calculation of tilt-dependent rate maps. The average firing rate was calculated for directions of the gravity vector (***a^G^***, in head coordinates) falling within 20° (green circle) of a series of points evenly distributed over a sphere (black dots). (H) Tilt-dependent rate maps for 4 example A^G^-units (top) and 4 example **Ω**-units (bottom). Dashed circles represent the equator (90° head tilt). (I) Boxplot of the CV of firing rate values in tilt-dependent rate maps for **Ω**-units (*n* = 6) and A^G^-units (*n* = 12). \*\**p* < 0.01.

In comparison to **Ω**-units, A^G^-units exhibited minimal rotational sensitivities (0.03 ± 0.02 °^−1^, *n* = 7, *p* = 1.2e−3; Fig. 3E). The tuning of A^G^-units was examined by computing their average firing rate as a function of head tilt (i.e. the orientation of the gravity vector ***a^G^*** in head coordinates; Fig. 3G). The resulting plots confirmed that A^G^-units were strongly modulated by head tilt (Fig. 3H), contrarily to **Ω**-units (CV higher for A^G^-units than for **Ω**-units: 2.28 vs. 0.29, *p* = 4.7e−3; Fig. 3I). A^G^-units were classified as mossy fibers (*n* = 1), Golgi cells (*n* = 3) and Purkinje cells (*n* = 8). Overall, these data show that a fraction (20%) of caudal cerebellar units displayed selective tuning to either head angular velocity or head tilt; most of our granular layer units (6/8) belonged to these categories.

### A population of units displays tilt-dependent rotational tuning

As shown above, most units displayed a mixed sensitivity to rotational and gravitational information and were not classified as **Ω**-units or A^G^-units. A direct examination of inertio-temporal receptive fields for different head orientations showed that the rotational sensitivity of these units was indeed often highly tilt-dependent.Fig. 4–F shows two example units for which the apparent sensitivity to ***ω**_z_* increased (Fig. 4A) or even reversed (Fig. 4D) for positive vs. negative values of 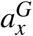(nose down vs. up), as confirmed by the slope of firing rate vs. ***ω**_z_* regressions for positive vs negative values of 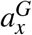(Fig. 4B,E)and by plotting the firing rate as a function of *ω_z_* and head elevation angle at the optimal lag (Fig. 4C,F). We therefore calculated a stability index σ quantifying the effect of 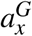(*σ_x_*) or 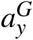 (*σ_y_*) on the direction of rotational sensitivity vectors over a given lag range (a maximal value of 1 indicating identical directions for opposite head orientations; see Methods). This index was lower for non **Ω**-units (*σ_x_*= 0.36 ± 0.32 and *σ_y_* = 0.41 ± 0.29, *n* = 60) than for **Ω**-units (*σ_x_*= 0.73 ± 0.13 and *σ*_y_ = 0.88 ± 0.06, *n* = 6; *p* = 2.7e−3 and *p* = 1.1e−4, respectively; Fig. 4G,H), confirming that the rotational tuning of this population was head tilt-dependent.

**Figure 4.**
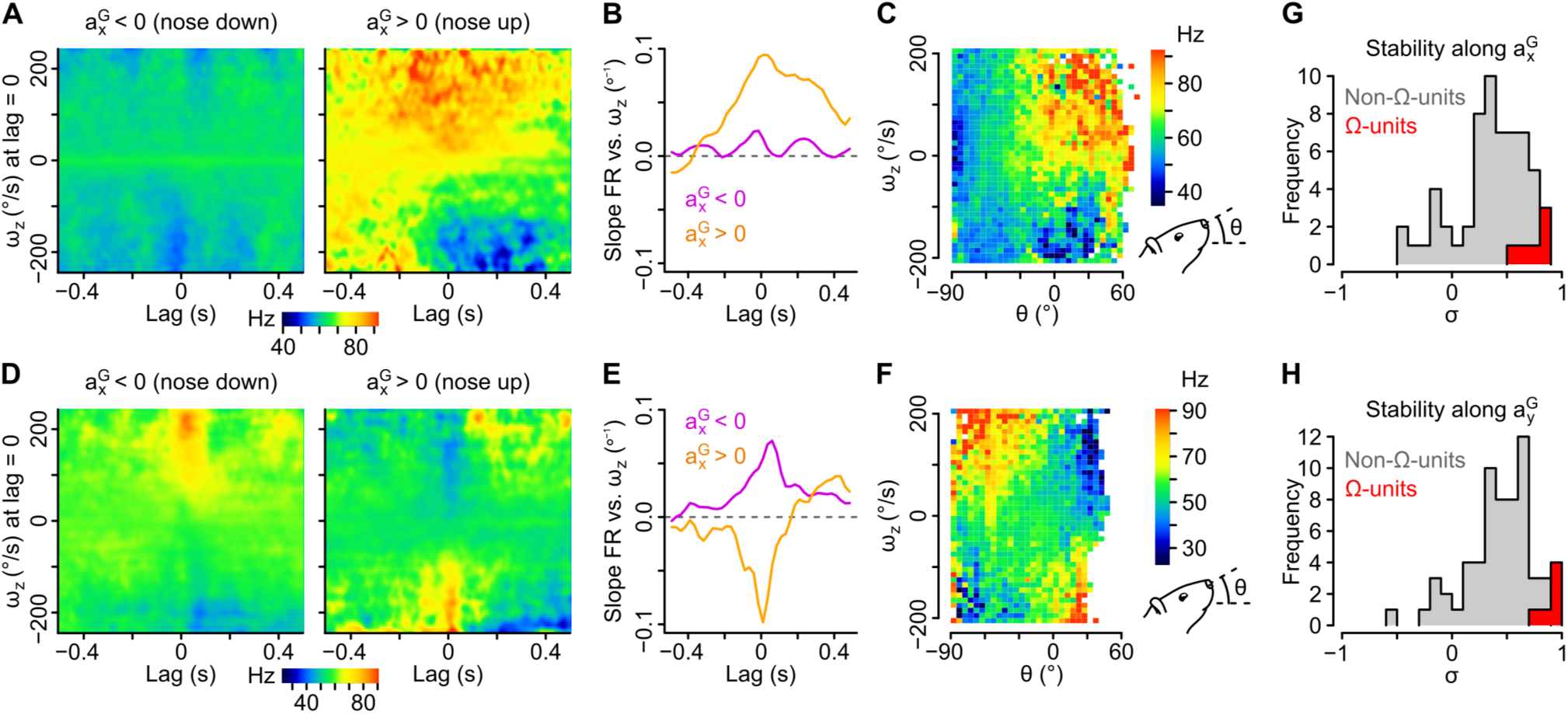
Tilt-dependence of rotational sensitivity in units with mixed gravitational and rotational sensitivity. (A–F) Example units exhibiting a pitch tilt-dependent modulation of their apparent sensitivity to yaw velocity (*ω_z_*, measured in the sensor’s frame). In one unit (A–C), ***ω**_z_* sensitivity is visible for nose up orientations only (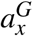> 0). In the other unit (D–F), ***ω**_z_*, sensitivity reverses for nose up vs. down orientations (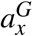> 0 vs. 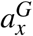 < 0). A & D: Inertio-temporal receptive fields for ***ω**_z_*, for nose up vs. down orientations. B & E: Slope of the firing rate vs. ***ω**_z_* linear regression (calculated from the receptive fields in A and D), plotted for different lag values in the nose up and nose down orientations. C & F: Histogram showing the average firing rate (color coded) as a function of ***ω**_z_* and of the head’s pitch angle (*θ*). (G–H) Histograms of the stability index calculated for positive vs. negative values of 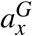 (*G*) and for positive vs. negative values of 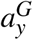(*H*). The stability index was used to quantify the influence of head tilt on the direction of rotational sensitivity over a given lag range. Values close to 1 (resp. −1) denote a weak (resp. strong) influence of head tilt on the direction of rotational sensitivity. Histograms for non-**Ω**-units with significant rotational sensitivity (*n* = 60 units) are colored in gray and histograms for **Ω**-units (*n* = 6 units) are colored in red.

### A fraction of cerebellar units encodes head rotations in a gravity-centered reference frame

The above data suggest that some units employed a rotational coding scheme that takes into account the direction of gravity, raising the possibility that some of them might encode head rotations in a reference frame aligned with the earth-vertical direction. To explore this, we computed rotational sensitivity vectors (***ω_opt_***) for all head tilt angles explored by the animal, using angular velocity signals expressed either in an internal (head-bound), or in an external (earth-bound), reference frame (Fig. 5B; see Methods). We reasoned that ***ω_opt_*** vectors of units encoding rotations in one reference frame (internal of external) should remain aligned (tilt-independent) only when calculated in this particular frame. As a means to visualize the effect of head tilt on these vectors, we plotted them on a sphere at the coordinates corresponding to the head tilt (i.e. the coordinates of gravity in the head reference frame) for which they were calculated (Fig. 5A).Fig. 5C,D shows two example units with different tuning properties. In the left unit (Fig. 5C), ***ω_opt_*** vectors appeared more consistently aligned when calculated in external (vs. internal) coordinates, while the opposite was observed for the right unit (Fig. 5D; see Movie 2 for a 3D version of these plots). This was confirmed by examining the collinearity of ω vectors as a function of the angular distance between their loalization on the sphere: collinearity decreased with angular distance for internal but not external ***ω_opt_*** vectors in the left unit (Fig. 5E), while it decreased for external but not internal ***ω_opt_*** vectors in the right unit (Fig. 5F). Other units exhibited no clear preferred orientation for internal vs. external ***ω_opt_*** vectors and collinearity curves that decayed similarly for the two reference frames (Figure 5–figure supplement 1A,B). When plotted for all units, the difference between collinearity curves of external and internal ***ω_opt_*** vectors revealed a continuum of properties (Fig. 5G). Units exhibiting a stronger tuning toward an external or internal coding scheme were isolated by setting a cutoff (difference between external and internal collinearity > 0.5 or < −0.5 for angular distances ranging 80−100°). These units all corresponded to putative Purkinje cells. The preferred rotation axes of external-coding cells did not appear to cluster around specific directions (Fig. 5H).

**Figure 5.**
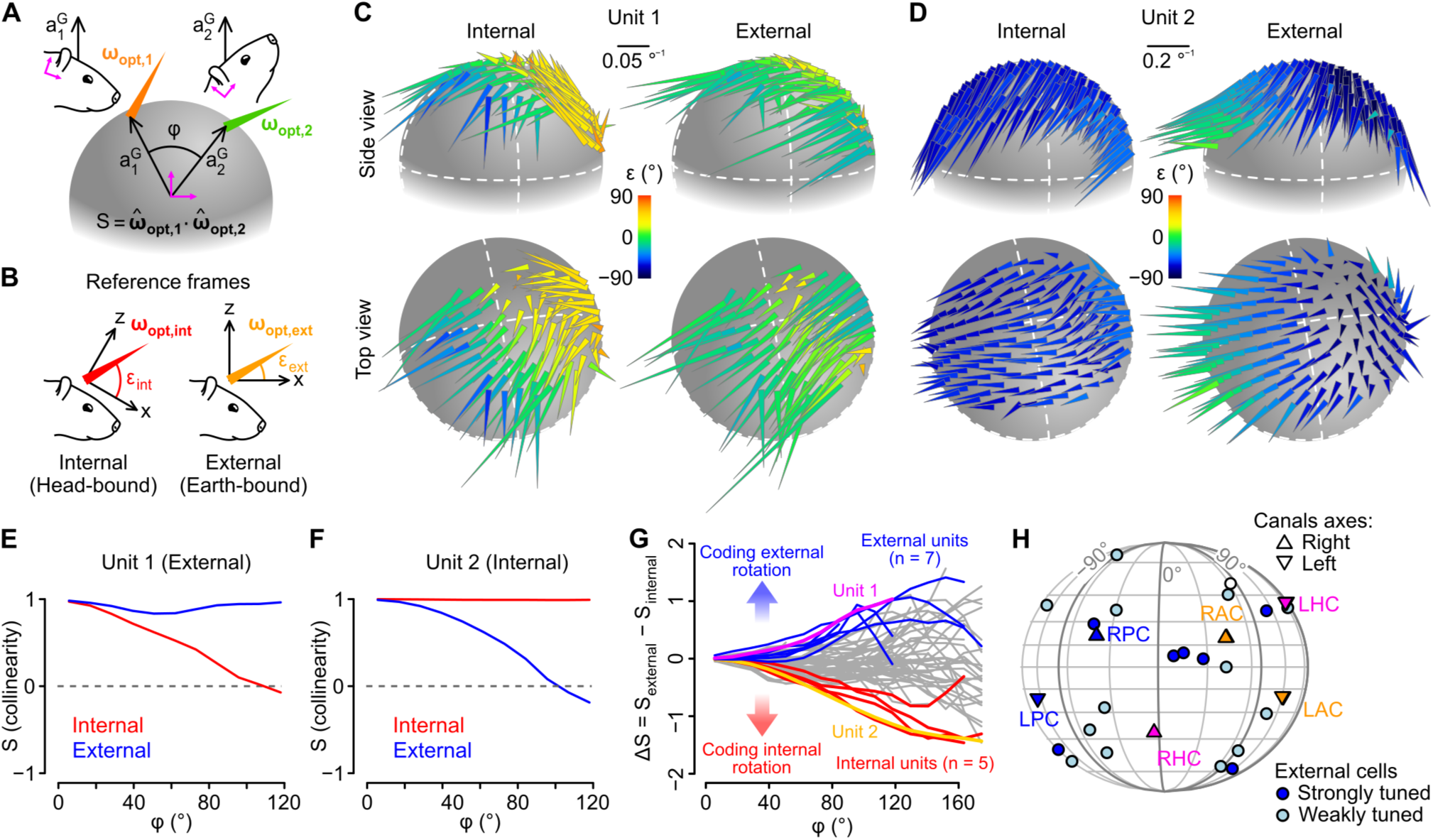
Different cerebellar units encode head rotations in a head-bound or earth-bound reference frame. (A) Method used to examine the influence of head tilt on rotational sensitivity. Optimal sensitivity vectors (***ω_opt_***) were calculated for different orientations of the gravity vector (***a^G^***) in head coordinates (***φ***: angular distance between different ***a^G^*** orientations). Collinearity (*S*) was assessed by computing the dot product of normalized sensitivity vectors. (B) ***ω_opt_*** vectors were calculated using internal (head-bound) or external (earth-bound) angular velocity values. ***ε***: angle of elevation relative to the (*x, y*) plane of the corresponding reference frame. (C–D) ***ω_opt_*** vectors of two examples units, calculated using internal (left) or external (right) angular velocity, positioned at locations corresponding to orientations of ***a^G^*** (in head coordinates) for which they were calculated, and color-coded according to their angle of elevation ***ε***. (E–F) Collinearity of externally- (blue) and internally-referenced (red) sensitivity vectors vs. angular distance for the two units shown in *C* and *D*. (G) Difference between external and internal collinearity curves (Δ*S*) for all units with significant rotational sensitivity (*n* = 66). Units with a strong external or internal tuning Δ*S* > 0.5 or Δ*S* < −0.5 for ***φ*** in the 80−100° range) were highlighted in blue (*n* = 7) and red (*n* = 5), respectively. The two units shown in *C* and *E* were highlighted in purple (unit 1) and orange (unit 2). (H) Direction of external sensitivity vectors for the 7 units highlighted in blue in *G* (dark blue circles), and for 12 units with a weaker external tuning (Δ*S* > 0.1 for ***φ*** in the 80−100° range, light blue circles), plotted on a pseudocylindrical projection. The excitatory direction of rotation of semi-circular canals is indicated as in Fig. 3F).

Our description of the units’ sensitivity to rotational and gravitational information hypothesizes a linear tuning to tilt-dependent rotation axes. To determine whether this description is as good as our model-free resampling approach at explaining firing rate modulations, we compared *R*^2^ values calculated from tilt-dependent linear fits with the ones obtained by resampling. The comparison showed that both values were similar (*p* = 0.24, *n* = 86 units; Figure 5–figure supplement 1C), supporting the idea that the sensitivity of most units to head movements can be described as tilt-dependent linear responses to angular velocity signals.

## Discussion

Our findings reveal that the caudal cerebellar vermis hosts gravitationally-polarized representations of head rotations in freely moving rats. These representations corresponded to Purkinje cell receptive fields encoding head rotations about 3D axes anchored to the direction of gravity. This type of head tilt-independent rotational sensitivity requires a complex re-encoding of head-centered sensory cues and might subserve downstream computations such as the encoding of head direction in the earth-horizontal plane.

Head tilt affects the correspondence between head-bound and earth-bound rotations (see Movie 3). A cell tuned to rotations about an earth-bound axis shall thus exhibit a tilt-dependent sensitivity to head-bound rotations. We found indeed Purkinje cells with a tilt-dependent sensitivity to “internal” rotations (as measured by semi-circular canals) but which exhibited a more consistent sensitivity to “external” rotations (documenting changes in heading and head attitude). This population of cells provides a neuronal signal that could be used in forebrain structures to generate tilt-dependent rotational sensitivity (Laurens et al., 2016) and to update externally-referenced head direction signals through temporal integration (Taube et al., 2013; Finkelstein et al., 2016; Wilson et al., 2016).

The rotational sensitivity of caudal cerebellar units was maximal for positive lags relative to angular velocity suggesting that it was driven by sensory cues rather than motor commands. We found no evidence for a crucial role of visual inputs, but instead found differences in the sensitivity to active vs. passive movements. This result is reminiscent of the lower sensitivity of certain vestibular nuclear neurons to active vs. passive movements (Cullen & Roy, 2004), and might reflect a similar mechanism of attenuation of self-generated inputs. Alternatively, self-generated movements occurred at higher frequencies than our passive movements (⋍ Hz) and may thus yield smaller Purkinje cell modulations (Yakusheva et al., 2008). Further studies are required to identify the cause of differences of neuronal sensitivity to active vs. passive movements.

We also identified units tuned to either head tilt or head rotations. Most of our granular layer units belonged to these categories, thus potentially reflecting the activity of otolithic and semi-circular mossy fibers or Golgi cells directly driven by these fibers. A fraction of head tilt-selective units (*n* = 8) was identified as Purkinje cells, and might correspond to previously identified static roll-tilt Purkinje cells (Yakhnitsa & Barmack, 2006), or to tilt-selective Purkinje cells dynamically extracting head tilt information through multisensory integration (Laurens et al., 2013).

Finally, we found that the instantaneous firing rate of many units exhibited strong fluctuations even during complete immobility. As a result, the instantaneous firing rate was only loosely connected to the rapid component of head inertial signals, consistent with a coding of information at the level of Purkinje cell populations (Herzfeld et al., 2015), read through the summation of conergent Purkinje cell inputs in vestibular nuclear neurons. We found indeed that neighboring cells exhibited similar sensitivities to head movements, consistent with a microzonal organization of the cerebellar cortex (Dean et al., 2010).

Deciphering how Purkinje cell receptive fields may be tuned to rotations about fixed directions relative to gravity is a complex topic beyond the scope of this work. One challenge is to explain how rotational sensitivity is gated by gravity. Such an operation might be performed in the granular layer, by first computing an estimate of gravitational acceleration and then combining it with rotational (vestibular, visual or proprioceptive) informations. We found that the low-frequency (<2 Hz) component of acceleration during free movements mainly contains gravitational information. The granular layer of the caudal vermis contains a high amount of unipolar brush cells (excitatory interneurons intercalated between mossy fibers and granule cells) which may smooth otolithic signals over hundreds of milliseconds (van Dorp & De Zeeuw, 2014; Borges-Merjane & Trussell, 2015; Zampini et al., 2016) and thus provide a proxy of gravitational signals to granule cells. Granule cells receiving convergent inputs carrying gravitational and rotational information could then operate as coincidence detectors (Chadderton et al., 2004; Chabrol et al., 2015) and signal the occurrence of specific combinations of rotation and head tilt to Purkinje cells.

Purkinje cell receptive fields are shaped by the activity of climbing fibers which determine the sensitivity to subsets of granule cells (parallel fiber) afferents (e.g. Dean et al., 2010). Vestibular climbing fibers emanate from inferior olive neurons of the *β* nucleus, which are controlled by dorsal Y-group and parasolitary nucleus afferents. These nuclei carry rotational and low pass filtered otolithic signals but are also under the influence of vestibulo-cerebellar Purkinje (Barmack & Yakhnitsa, 2000; Barmack, 2003; Wylie et al., 1994). Therefore, the teaching signal sent to Purkinje cells by way of olivary neurons shall result from a complex interplay between external afferents and the action of Purkinje cells themselves, leading to the observed tilt-dependent rotational sensitivity.

To conclude, as emphasized previously (Green & Angelaki, 2007), the recoding of head-bound sensory cues into earth-bound kinematic signals involves non-linear operations. The detailed mechanisms underlying operations leading to the emergence of rotational receptive fields anchored to gravity in the caudal cerebellar vermis remain to be elucidated.

## Materials and Methods

Full details of the procedures are provided in the Supplemental Experimental Procedures. Details of statistical tests are provided in the Supplemental Experimental Procedures. All *p*-values were obtained with an unpaired Wilcoxon test. All mean values are given with the standard deviation.

## Author Contributions

GPD, MT and CL designed the experiments. MT performed the experiments. GPD, MT, BG and CL designed and performed the analysis. GPD, MT, BG and CL wrote the manuscript.

## Acknowledgements

We thank J. Laurens, D. Bennequin, A. Afgoustidis and D. Popa for helpful discussions on the project, and L. Rondi-Reig and C. Rochefort for initial discussions. We thank M. Pasquet, Y. Cabirou and G. Paresys for technical assistance. This work was supported by grants from France’s Agence Nationale de la Recherche (ANR) to C.L. (ANR-12-BSV4-0027) and to B.G. (ANR-15- CE37-0007), and has received support under the program “Investissements d’Avenir” launched by the French Government and implemented by the ANR, with the references ANR-10-LABX-54 MEMO LIFE and ANR-11-IDEX-0001-02 PSL* Research University.

